# Cholesterol Sequestration by Xenon Leads to Lipid Raft Destabilization

**DOI:** 10.1101/2020.05.04.077727

**Authors:** A.D. Reyes-Figueroa, Mikko Karttunen, J.C. Ruiz-Suárez

## Abstract

Combined coarse-grained (CG) and atomistic molecular dynamics (MD) simulations were performed to study the interactions of xenon with model lipid rafts consisting of 1,2-dipalmitoyl-sn-glycero-3-phosphocholine (DPPC), 1,2-dilauroyl-sn-glycero-3-phosphocholine (DLPC) and cholesterol (Chol). At a concentration of 2 Xe/lipid we observed an unexpected result: Spontaneous nucleation of Xe nanoclusters which then rapidly plunged into the bilayer. In this process Chol, essential for raft stabilization, was pulled out from the raft into the hydrophobic zone. When concentration was further increased (3 Xe/lipid), the clusters disrupted both the membrane and raft. We computed the radial distribution functions, pair-wise potentials, second virial coefficients and Schlitter entropy to scrutinize the nature of the interactions. Our findings suggest that the well-known anaesthetic effect of Xe could be mediated by sequestration of Chol, which, in turn, compromises the stability of rafts where specialized proteins needed to produce the nervous signal are anchored.

---

Despite being used in countless hospitals daily, the molecular mechanism(s) that control general anesthesia remain unknown. The range of molecules capable of producing it is very broad, the most famous being propofol, halogenated agents, laughing gas (nitrous oxide) and, perhaps the most stereotypical one, xenon [1]. Although it remains elusive why such diverse agents produce anesthesia, the pioneering works of Meyer [2] and Overton [3], thoroughly studied by Heimburg *et al.* [4–6], have shown that general anesthesia may be directly related to the partitioning coefficient. Current hypotheses include the alteration of membrane properties and, consequently, the functions of proteins via changes in the lateral pressure profile and the membrane potential [1, 7–12].

The proposed action mechanism for Xe, according to a canonical report [13], is that it contends for binding at the site where the neurotransmitter glycine is needed to activate the N-methyl-D-aspartate receptor (NMDA). This, however, comes with at least two dilemmas: 1) How does an inert atom *selectively* block the NMDA calcium channel [14] and hence the neuronal signal, and 2) is it possible that there are several mechanisms for general anaesthetics?

The authors that first proposed Xe as an antagonist agent suggest a solution: Xe binds at the GluN1 subunit of the NMDA receptor [13], precisely in the aromatic ring of the Phe-758 residue [15]. A recent report goes even further [16]: The binding energy of Xe with this aromatic ring is 11.3kJ/mol and the distance to the plane of the six carbons composing such a ring between 3.5 and 5Å.

Despite the above suggestions [13, 16] important controversies remain: 1) If Xe enters into the protein to quay a few Ångstroms away from the aromatic ring, how does it manage to suppress the hydrogen-bond donors on the protein receptor where glycine has its specific action? 2) The proposed antagonist capability of Xe is at odds with its role as a contrast augmenter in protein crystallography, precisely because the structures of proteins with and without Xe are highly isomorphous to each other, i.e., proteins suffer only marginal changes [17, 18].

What if instead of acting directly on a protein receptor or attractive cavities, Xe atoms enter into the membrane, change its structural stability and, as a consequence, the functions of the receptors moored in there become perturbed? This idea is not new [7] and studies of membrane structural changes produced by Xe have been performed [19–23]. However, as far as we know, the effects of Xe on lipid rafts have been studied only marginally. The most notable is perhaps the diffraction study of Weinrich and Worcester, albeit with a different conclusion [24].

Rafts are small, heterogeneous, highly dynamic, and sterol- and sphingolipid-enriched membrane domains that are able to compartmentalize receptors and membrane-proximal effectors [25, 26]. They are, at physiological temperatures, in the liquid-ordered phase with intermediate characteristics between gel and the liquid-disordered phase, in coexistence with the rest of the membrane that is in the liquid-disordered phase [27]. The high affinity of some proteins to these microdomains facilitates the formation of complexes and the activation of specific signaling pathways [28], and it has been found that unspecific binding in lipid membranes modifies lipid rafts [29, 30]. This has been shown to be the case, for instance, in virus fusion [31], immune modulation [32], cancer [33], Alzheimer disease [34], endocytosis [35], T-cell activation [36] and neurotransmitter signaling [37].

We used a combination of Martini coarse-grained (CG) and atomistic MD simulations to study three concentrations (1,2 and 3 Xe/lipid) of Xe in lipid rafts composed of DPPC, DLPC and Chol both below and above the miscibility temperature at 295 and 323 K. Simulations with the Martini CG (version 2.2) parameter set “ new-rf” [38] were first performed. The system consisted of 238, 510 and 810 Chol, DPPC and DLPC, respectively, solvated with 13,448 water beads. The initial configuration was a random distribution of lipids obtained using CHARMM-GUI [39, 40]. This was followed by energy minimization using the steepest descents algorithm and a set of 1 ns pre-equilibration simulations using time steps (*τ*) of 2, 5, 10, 15 and 20 fs. The v-rescale thermostat [41] and the Berendsen barostat [42] at *T* = 295 K and 1 bar were used. The rest of the CG runs were done using the Parrinello-Rahman barostat [43]. Equilibration was completed by a high temperature cycle (400 K) for 100 ns and return back to 295 K. 100 ns, followed by a 10 ns simulation at 295 K. The production runs were for 7 *μs* using *τ* = 10 fs. The lipid raft formed spontaneously around 2.5 *μs*, snapshots are shown in Fig. S1.

The last structure of this CG simulation was converted to an atomistic model using the method of Wassenaar *et al.* [44]. The GROMOS 43A1-S3 force field optimized by Chiu et *al.* [45] was used with the DLPC parameters from Ref. [46]. The Simple Point Charge [47] model was used for water. Following previous works for Xe [20, 21], the Lennard-Jones parameters *e* =1.8370 and *σ* = 0.4100 were employed. As with CG simulations, energy minimization and equilibration (without the high temperature cycle) were performed. The final atomistic production runs were for 200 ns using the Nosé-Hoover thermostat [48, 49] and the Parrinello-Rahman barostat [43] with coupling constants of 5 fs and 5 ps, respectively. The long-range electrostatics interactions were calculated using the particle mesh Ewald algorithm [50] with a 1.0 nm cutoff, a Fourier spacing of 0.15 nm and a sixth order interpolation. All the simulations were performed using the GROMACS package version 5.1.5 [51].

The membrane in Fig. 1 corresponds to the case of 2 Xe/lipid at 295K (T = 323K is shown in Fig. S2), details in the figure caption. Figure S3, shows the order parameters of DPPC and DLPC, illustrating that DPPC is more ordered than DLPC.

**FIG. 1.**
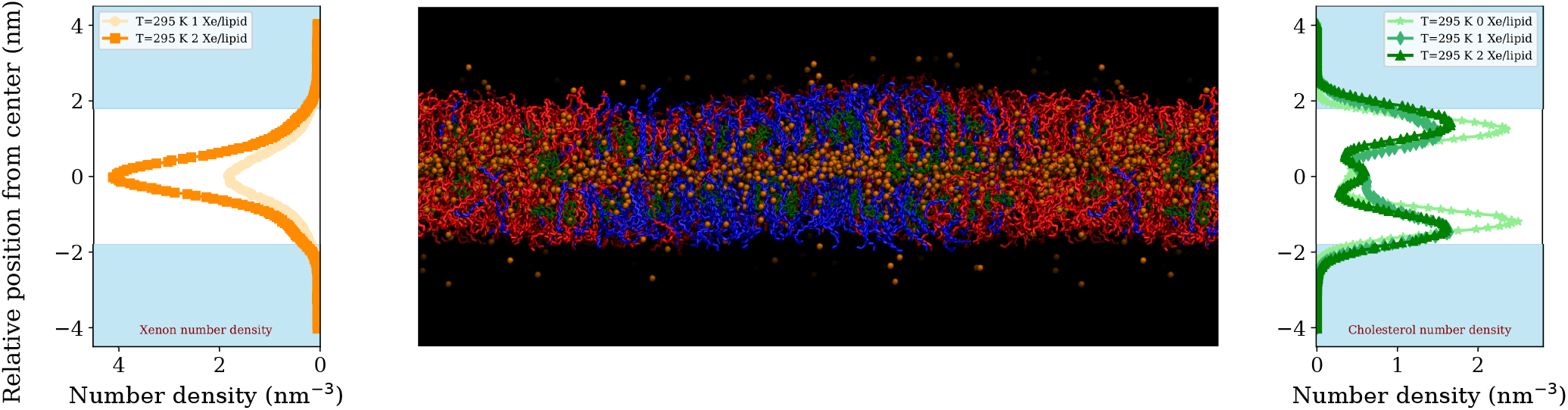
(color on line). A snapshot of a raft in a lipid membrane (2 Xe/lipid) at the end of the 200ns simulation at 295 K. DPPC, DLPC, cholesterol and Xe are depicted in blue, red, green and ochre colors, respectively. *Left:* The number density of Xe at two concentrations (1 and 2 Xe/lipid). Note the increasing density of Xe around the center of the bilayer (the white zone). *Right:* The number density for Chol at 0, 1 and 2 Xe/lipid. With no Xe, Chol density has two maxima around 1 nm from the center of the bilayer; at 2 Xe/lipid, the maximum density decreases and a peak emerges at the center.

Figure 2 shows the Xe-Xe pair distribution functions at the beginning (1 ns) and at the end (200 ns). At 1 ns the Xe atoms are homogeneously spread in the solution above the lipid membrane and at 200 ns the majority of them have migrated inside the lipid-tail zone. The radial distribution functions are qualitatively different, indicating that Xe is in different environments. We calculated the pair-wise Xe-Xe potential of mean force by inverting *g* (*r*), *v_PMF_*(*r*) = — *k_B_T* ln [*g*(*r*)], see the inset in Fig. 2. Clearly, the attractive part of the potential is much deeper when Xe is inside the membrane. The first minimum, when the Xe atoms are still in water, is around 0.41 nm, which corresponds to the value of *σ* in the Lennard-Jones potential. The second minimum (≃ 0.71 nm) corresponds to the situation where there is a water molecule, whose diameter is 0.3 nm, in between. At larger distances the potential goes to zero. While the first minimum in the deeper potential is around the same distance, the second one increases appreciably around 1 Å because the Xe atoms are separated now by lipid tails or C-H molecules (*σ* = 0.41 nm).

**FIG. 2.**
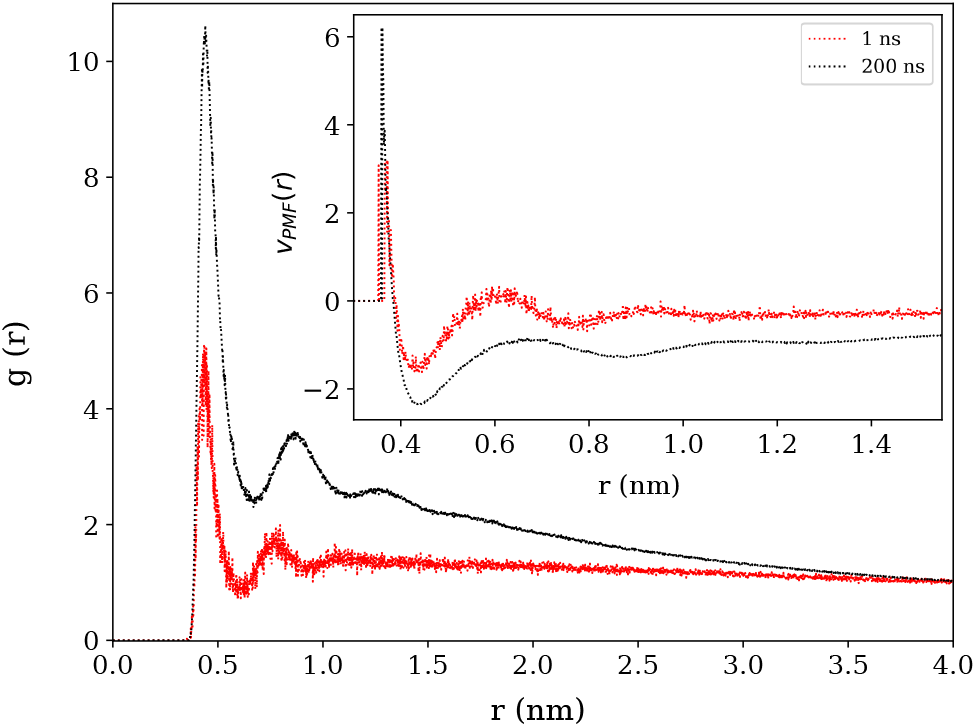
(color on line) The Xe-Xe pair distribution function *g*(*r*) at 295 K for 2 Xe/lipid at 1 and 200 ns. Inset:The potential of mean force calculated by inverting *g*(*r*), see text.

In order to examine the significance of the changes observed in the *g*(r)-functions, we evaluated the areas under the curves and then subtracted each value obtained at time *t* from the first value obtained at the beginning of the simulation,

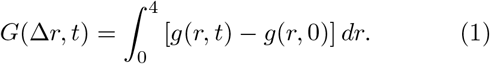

Time intervals of 5 ns were used to calculate *g*(*r,t*).

The main panel in Fig. 3 shows *G*(Δ*r, t*) for Chol-Xe at 295 and 323K. *G* increases, indicating that Chol molecules can not maintain their original positions and follow the Xe atoms towards the hydrophobic zone of the bilayer. The pair correlation functions for the Xe-Chol pairs is depicted in the inset of Fig. 3 at concentrations of 1 and 2 Xe/lipid. In addition, *G*(Δ*r, t*) in Fig. 4 shows the opposite behavior (decreasing) for Chol-DPPC at 1 and 2 Xe/lipid: *G* drops drastically, indicating that Chols separate from the DPPC lipids. The effects are intensified at 323 K (dashed lines). At 295 K, which is below the miscibility transition temperature of the lipid mixture in consideration [52], the raft is stable: G does not substantially change during the entire time of the simulation in the absence of Xe (solid red line).

**FIG. 3.**
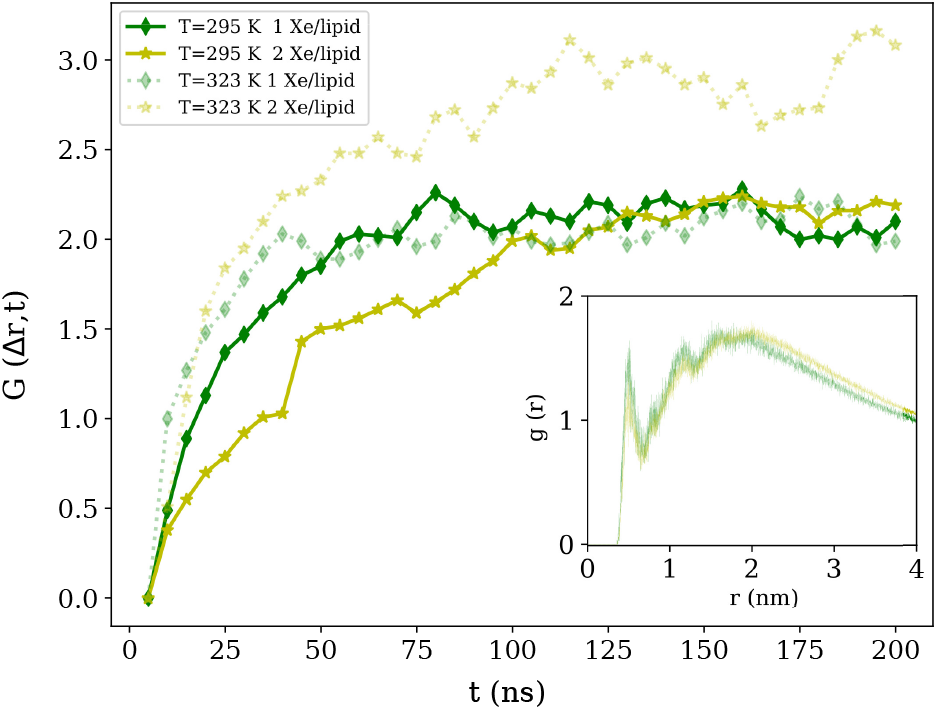
(color on line) Inset: The Xe-Chol pair distribution function at 295 K, where we observe a slight increment of *g*(*r*) (specifically at large distances) at 2 Xe/lipid, meaning that Chol is dragged inside the bilayer by the Xe atoms. The pair distribution functions are calculated for all times, but we show it only at the end of the 200 ns simulation. Main panel: *G*(Δ*r, t*) as a function of time for 1 and 2 Xe/lipid. The solid lines are for T=295 K while the dashed ones correspond to T=323 K. See text for details.

The pair correlation function for the Xe-Chol pair is depicted in the inset of Fig. 3. As the concentration is increased from 1 to 2 Xe/lipid, *g*(*r*) takes higher values for 2 Xe/lipid for *r* ≥ 2nm (i.e., the number of third nearest neighbors increases). This implies that Chol, first located in the immediate proximity of DPPC, detaches from it once Xe atoms penetrate the membrane. This is confirmed by the DPPC-Chol distribution function: *g*(*r*) reduces for *r* ≥ 1 nm (inset of Fig. 4).

**FIG. 4.**
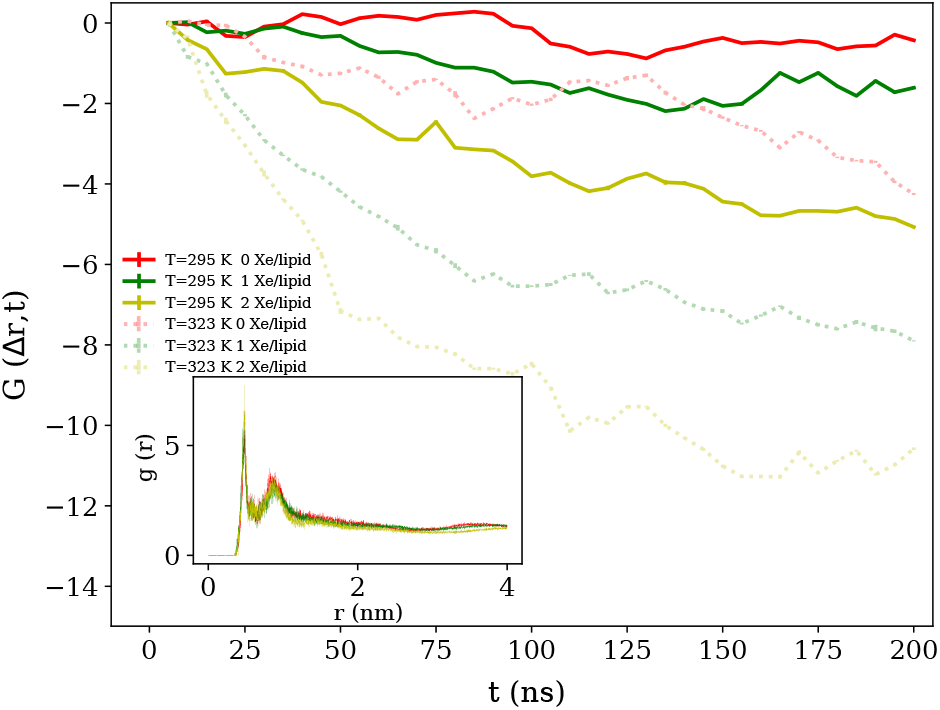
(color on line) DPPC-Chol pair distribution function at 295 K and 200 ns. At 2 Xe/lipid *g*(*r*) reduces, i.e. DPPC and Chol separate from each other. Inset: *G*(Δ*r, t*) as a function of time for 0, 1 and 2 Xe/lipid. The solid lines are for T=295 K while the dashed ones correspond to T=323 K. See text for details.

In order to quantify the nature and strength of the pairwise interactions, the second virial coefficients were evaluated using –2*π*∫ [*g*(*r*) — 1] *r*^2^*dr*, for four cases: Xe-Xe, Xe-Chol, Xe-DPPC, and DPPC-Chol, see Table I. The more negative the virial coefficient, the stronger the attraction. Thus, it is clear that Xe-Xe attraction is more intense in the hydrophobic membrane core: The virial coefficient changes from —18.34 in water to —55.08 nm^3^ inside the membrane. Furthermore, the virial coefficients of the Xe-Chol (DPPC-Chol) pairs are —35.36 (—44.98) nm^3^ at 1 Xe/lipid, but —43.74 (—24.56) nm^3^ at 2 Xe/lipid. Note also that the virial of the DPPC-Chol pair changes from −50 (0 Xe/lipid) to −24.56 nm^3^ (2 Xe/lipid). These changes in the virial coefficients confirm that both Xe and Chol sink into the hydrophobic core of the membrane.

**TABLE I.** Second virial coefficients (nm^3^) for Xe-Xe, Xe-Chol, Xe-DPPC, and DPPC-Chol at 295 K. The values were obtained at 200 ns, except those for Xe-Xe which correspond to 1 and 200 ns (in parenthesis).

| Xe/lipid | Xe-Xe | Xe-Chol | Xe-DPPC | DPPC-Chol |
| --- | --- | --- | --- | --- |
| 0 | - | - | - | -50 |
| 1 | - | -35.36 | -7.40 | -44.98 |
| 2 | -18.34 (-55.08) | -43.74 | -13.06 | -24.56 |

One could expect that the weakening of the interactions would lead to H-bond depletion and, therefore, to an increase in the entropy of the lipids. Chol molecules H-bond to the lipid headgroups. Thus, if the bonds become depleted, as suggested by the above results, the Chol molecules will be freer to move. The changes in the H-bonds are estimated in SI (see Figs. S4 and S5). Secondly, we calculated Schlitter’s entropy [53] to assess the conformational changes produced by Xe in lipid molecules using

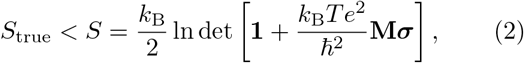

where **M** is the mass matrix, ***σ*** the covariance matrix with its elements defined as *σ_ij_* = 〈(*x_i_* — (〈*x_j_*〉)〉〈(*x_j_* — 〈*x_j_*〉)〉, *k_B_* is the Boltzmann constant, *T* the absolute temperature, *e* Euler’s number, and *ħ* is the Planck’s constant divided by 2*π*.

Translations and rotations around the center of mass of each molecule were removed by least-squares fitting of the trajectory configurations of the molecule during the calculation of the covariance matrix. In this form, the translational entropy can be excluded from the calculation but rotational entropy cannot be rigorously separated from internal motions of a flexible molecule. Then, an approximation of the entropy contributions of internal degrees of freedom (e.g., torsional angles) can be obtained using Cartesian coordinates and the above formula. *T*Δ*S* increases for each lipid, meaning that the disorder of the whole membrane increases substantially with the presence of Xe. The calculation also shows that the relative increase in entropy is the highest for cholesterol at 295 K, Table II. The importance of entropy has also been pointed out by Reigada and Sagues in connection with chloroform and its interactions with cholesterol containing membranes [54].

Figure S6 shows lateral and top views of the Xe-membrane system after 34 ns at the beginning of the atomistic simulation. Unexpectedly, before fully entering the membrane, Xe atoms cluster in the water phase, significantly above the rafts. Once the Xe clusters pervade into the hydrophobic lipid-tail zone, the absence of water molecules and therefore the suppression of the hydrophobic forces, makes the clusters spread within the bilayer, see the lateral view of the snapshot at 200 ns in Fig. 1. Upon increasing the concentration to 3 Xe/lipid, the system heads towards a catastrophic event, see Fig. S7 (295K) and Fig. S8 (323K); the accumulation of Xe and Chol within the bilayers leads to significant membrane expansion. This compromises the stability and integrity of the lipid rafts since Chol is a required component in them [52, 55].

**TABLE II.** Configurational entropy changes *T*Δ*S* in kJ/mol.

| Temperature | Xe/lipid | DPPC | Chol | DLPC | Total |
| --- | --- | --- | --- | --- | --- |
| 295 K | 1 | 10.13 | 6.58 | 11.67 | 28.38 |
| 295 K | 2 | 19.29 | 18.08 | 20.31 | 57.69 |
| 323 K | 1 | 14.00 | 15.93 | 14.44 | 44.37 |
| 323 K | 2 | 21.99 | 24.03 | 19.40 | 65.42 |

Summarizing, our MD simulations show that Xe atoms initially positioned outside the lipid membrane migrate to the lipid tail zone. Consequently, Chol molecules are pulled out from the liquid-ordered phase of the raft environment, probably because Xe atoms debilitate their interactions with the carbonyl groups and hydrophobic forces drive them inside. Our results are consistent with the scattering experiments of Weinrich and Worcester [24] who reported that the phase behavior of rafts becomes shifted toward the liquid-disordered phase upon addition of Xe, and with the simulations of Reigada using chloroform molecules in a liquid-ordered phase (DSPC/Chol bilayers) reporting an increase in membrane disorder when the drug molecules moved into the bilayer center [56]. At 2 Xe/lipid, the Xe atoms in water nucleate to form nanoclusters which then enter the membrane most notably through the rafts. At an even higher concentration (3 Xe/lipid) this outcome becomes further enhanced, and the membrane and the raft undergo an expansion due to the accumulation of Xe. Eventually this results in the loss of membrane integrity. Since the ion channels needed in neural signal propagation are believed to be located in the raft domains, our findings may offer a new explanation for general anaesthesia caused by Xe and other molecules.

This work has been supported by Conacyt, Mexico, Grant FC-1132. Computational Resources were provided by SharcNet/Compute Canada. ADRF acknowledges a scholarship from Conacyt, Mexico. MK thanks the Natural Sciences and Engineering Research Council of Canada (NSERC) and the Canada Research Chairs Program.

---

**Supplementary Figure S1.**
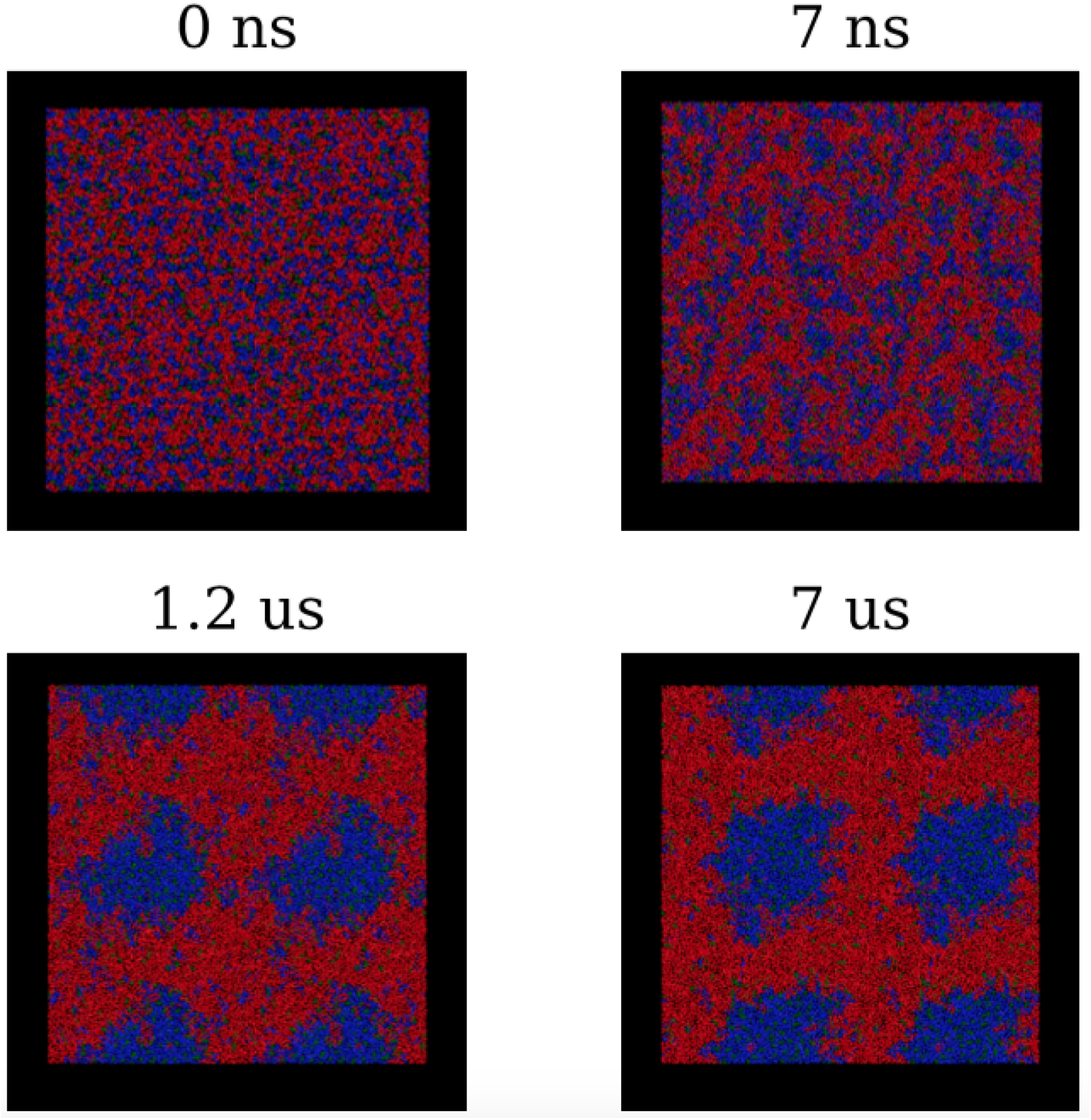
Formation of a lipid raft during a coarse-grained simulation. DPPC, DLPC, and cholesterol are depicted in blue, red, and green, respectively.

**Supplementary Figure S2.**
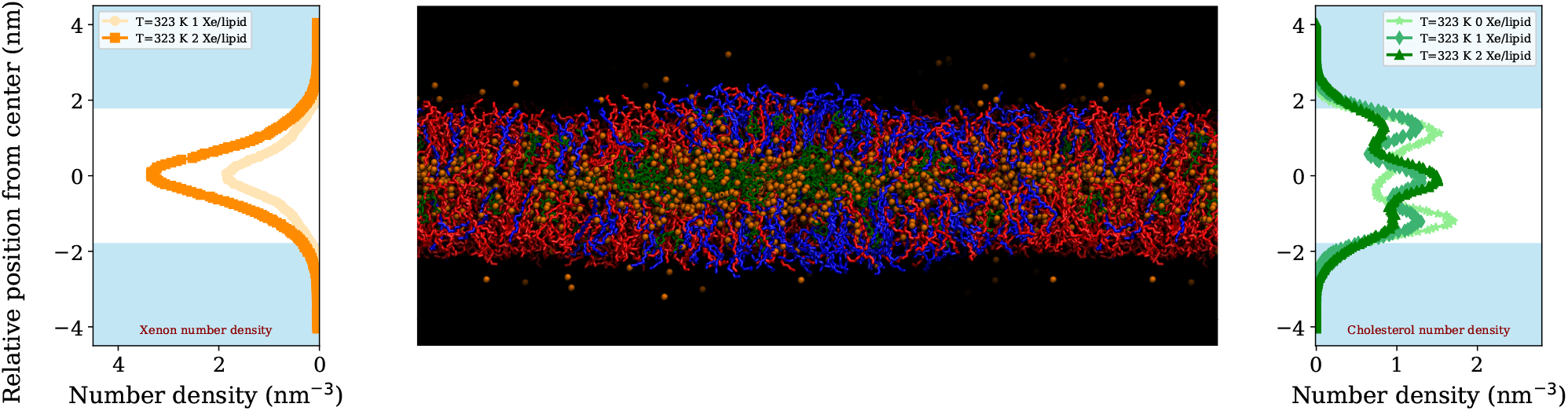
A snapshot of the lateral view of a lipid raft in a membrane at the end of the 200 ns simulation at 323 K. DPPC, DLPC, cholesterol and Xe are depicted with blue, red, green and ochre, respectively. The left panel illustrates the number density for Xe at two concentrations (1 and 2 Xe/lipid). Note the increasing density of Xe around the center of the bilayer indicated by the white zone. The right panel shows the number density for cholesterol at 0, 1 and 2 Xe/lipid. In the absence of Xe, the cholesterol density has two maxima around 1 nm from the center of the bilayer; at two xenon atoms per lipid, the maximum density decreases and a large peak shows up in the center. The snapshot of the membrane corresponds to 2 Xe/lipid.

**Supplementary Figure S3.**
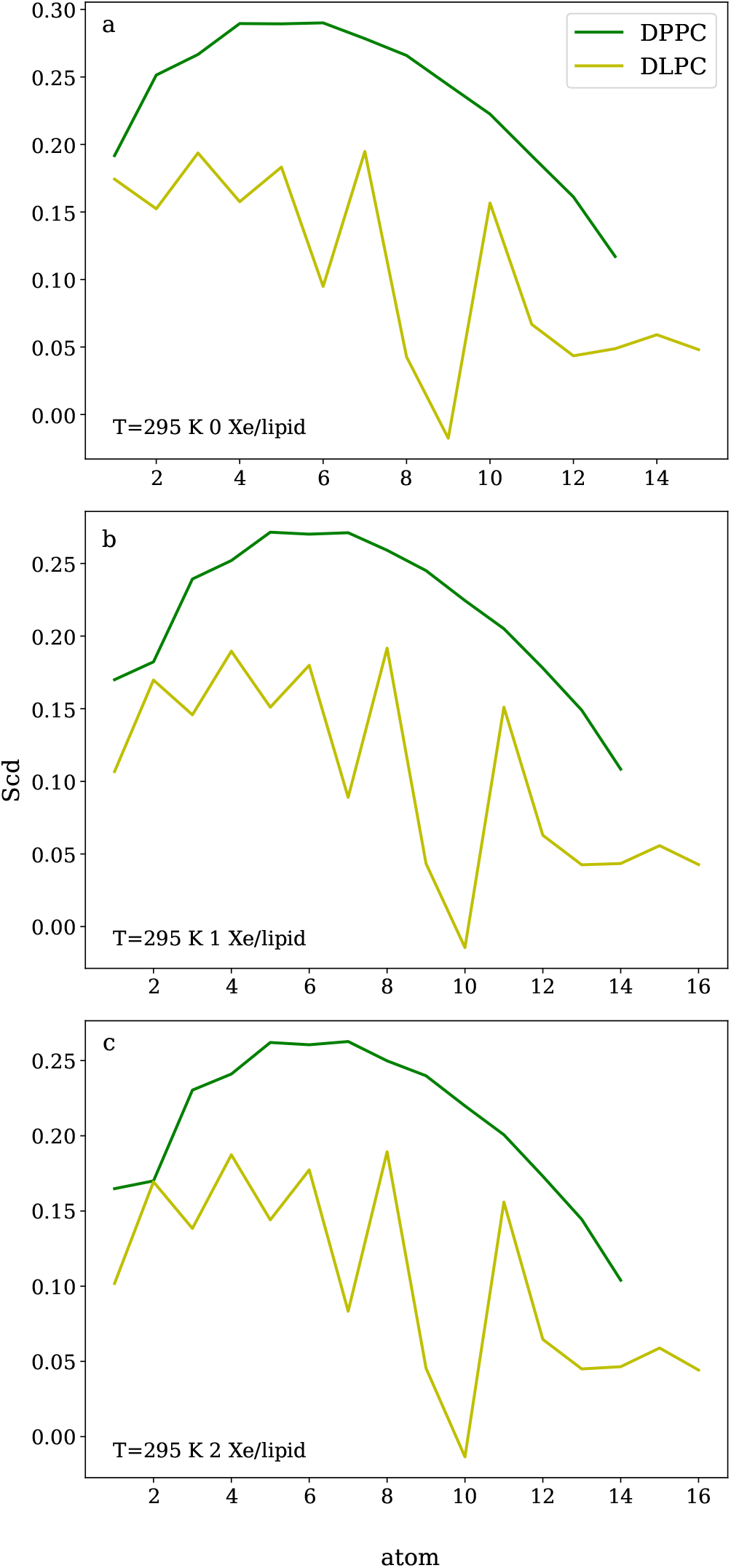
Order parameters for DPPC and DLPC, for 0, 1 and 2 Xe atoms/lipid at 295 K. Clearly, DPPC is more ordered.

**Supplementary Figure S4.**
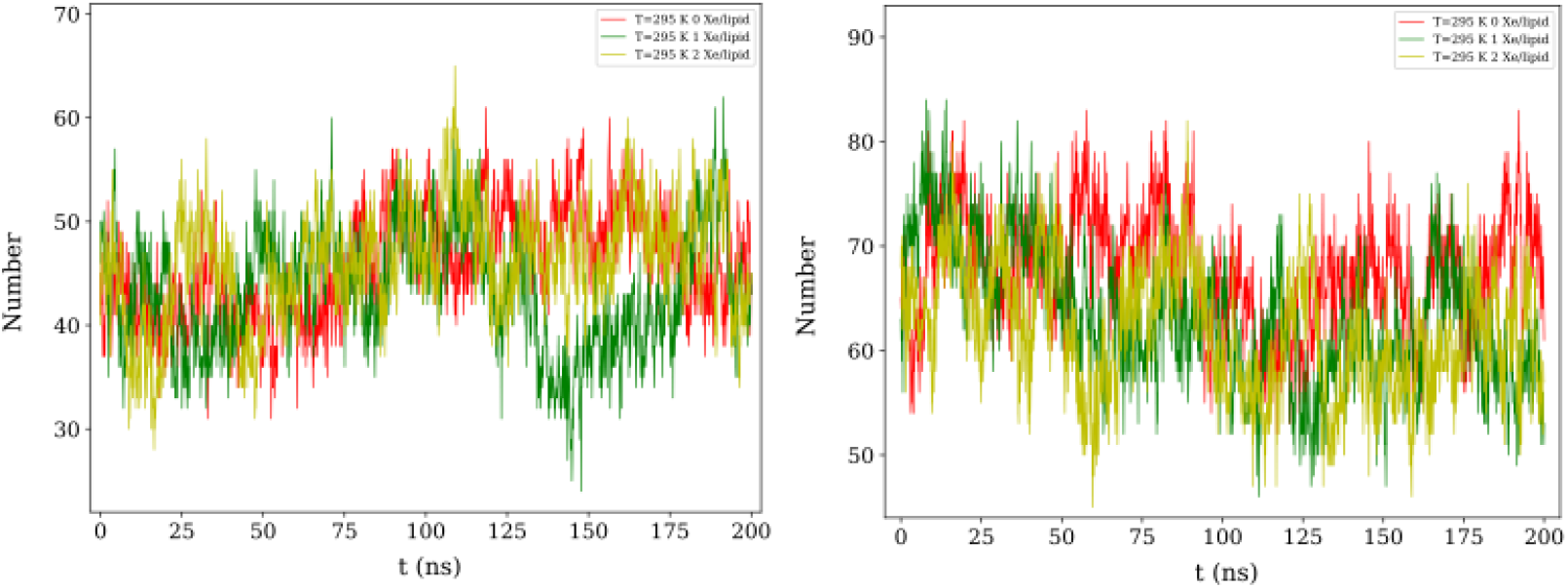
Number of hydrogen bonds between DLPC/DPPC and cholesterol vs time at 295 K, for 0, 1 and 2 Xe/lipid. Note that while in the non-raft zone (DLPC) the number of HBs remains constant during the simulation, in the raft zone (DPPC) there is a slight but clear reduction in the number of bonds at both Xe concentrations.

**Supplementary Figure S5.**
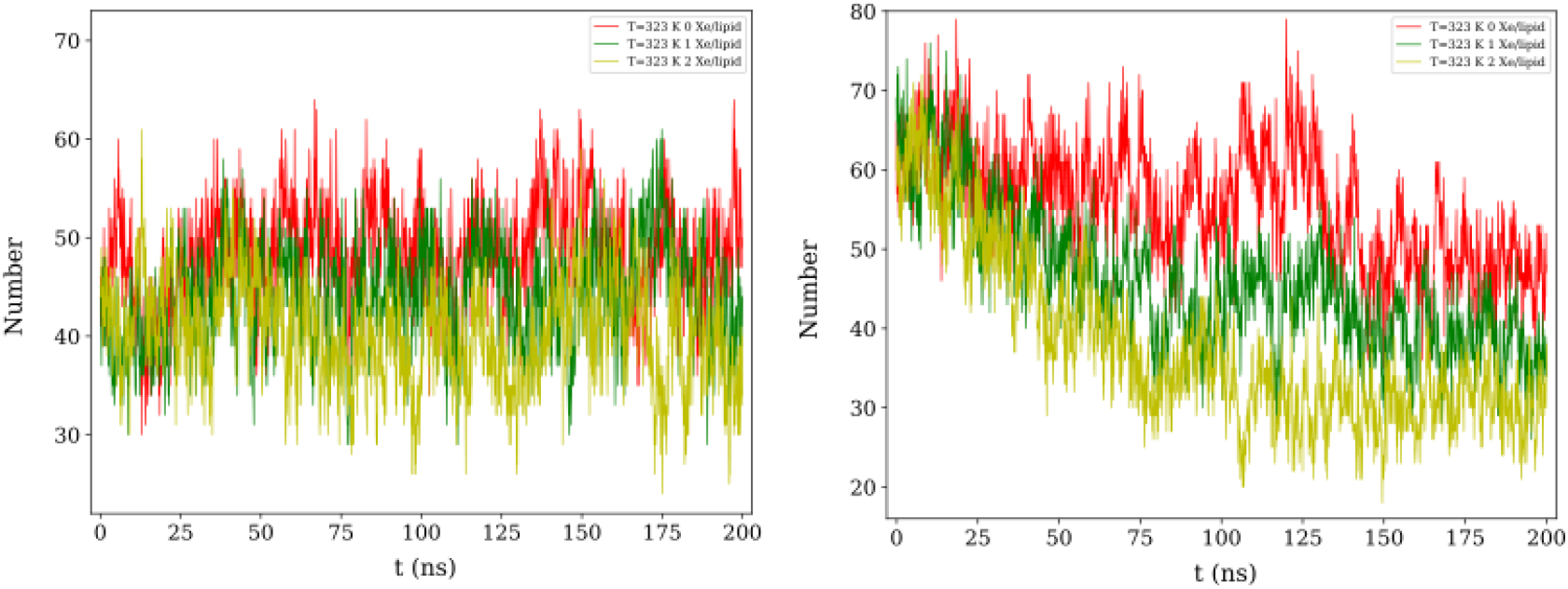
Number of hydrogen bonds (HB) between DLPC/DPPC and cholesterol vs time for 0, 1 and 2 Xe/lipid. In the non-raft zone (DLPC) the number of HB remains constant during the simulation, although the number is slightly higher in the absence of Xe. In the raft zone, however, even in the absence of Xe the number of HB reduces, indicating the gradual decomposition of the raft at this temperature; as indicated in the main text, it is above the miscibility temperature. At both Xe concentrations, the reduction of the HB drops illustrating the strong effect of Xe atoms.

**Supplementary Figure S6.**
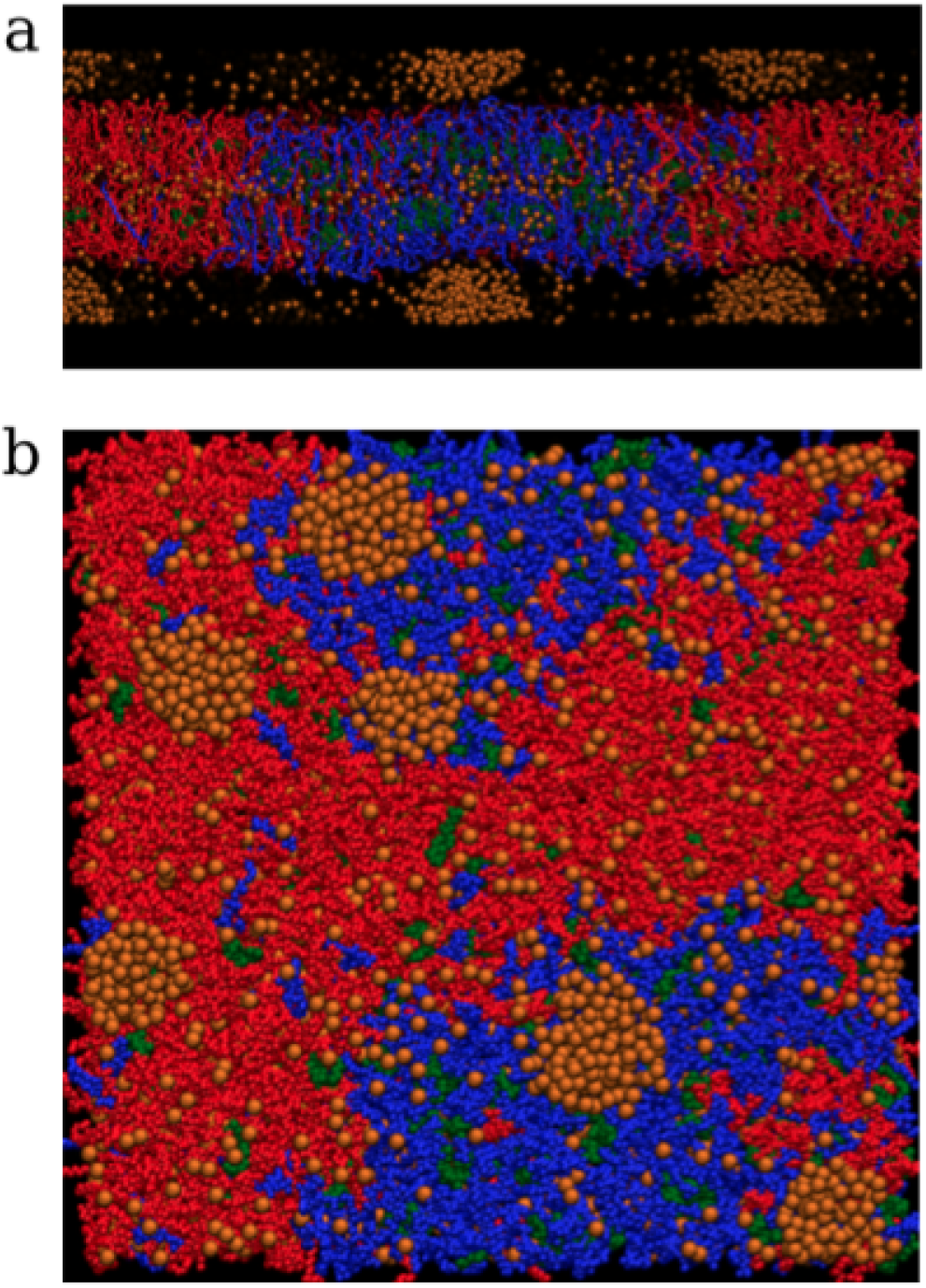
(a) Lateral view of the lipid membrane and the raft. As in Figure 1, cholesterol is marked in green. Note that at 2 Xe/lipid, Xe aggregates in nanoclusters before partitioning into the interior of the membrane. (b) Top view of the membrane/raft system.

**Supplementary Figure S7.**
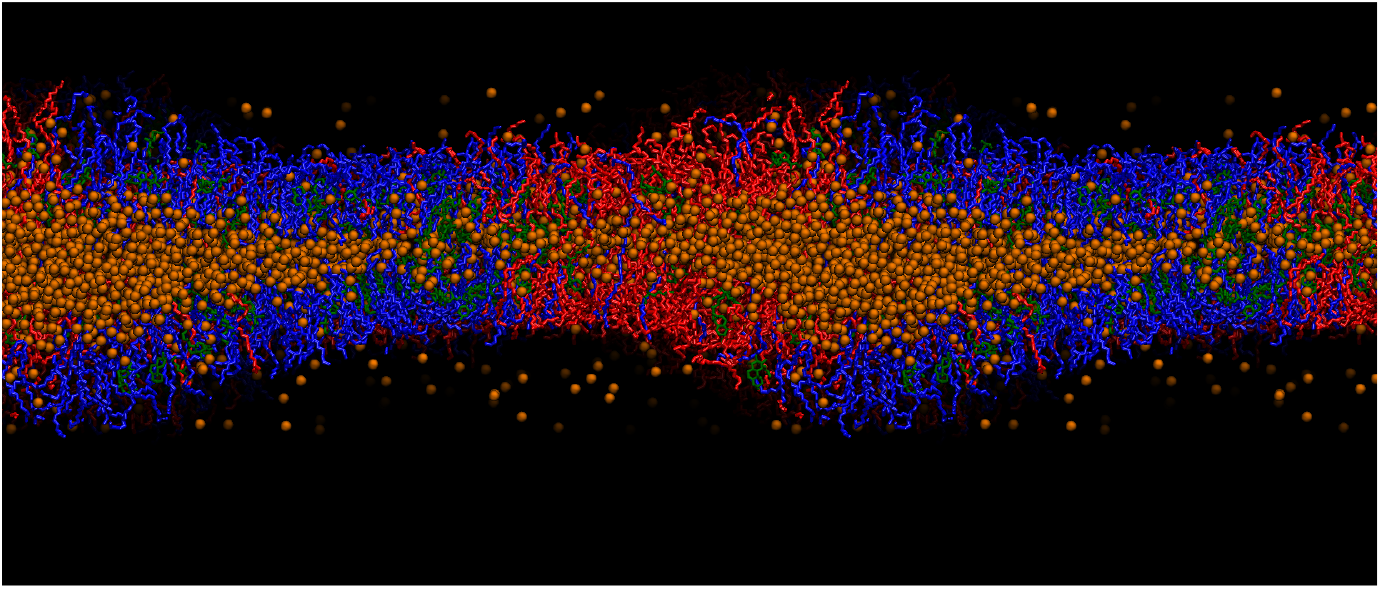
At the concentration of 3 Xe/lipid, the bilayer expands due to the copious accumulation of Xe atoms and cholesterol (200 ns). The temperature is 295 K.

**Supplementary Figure S8.**
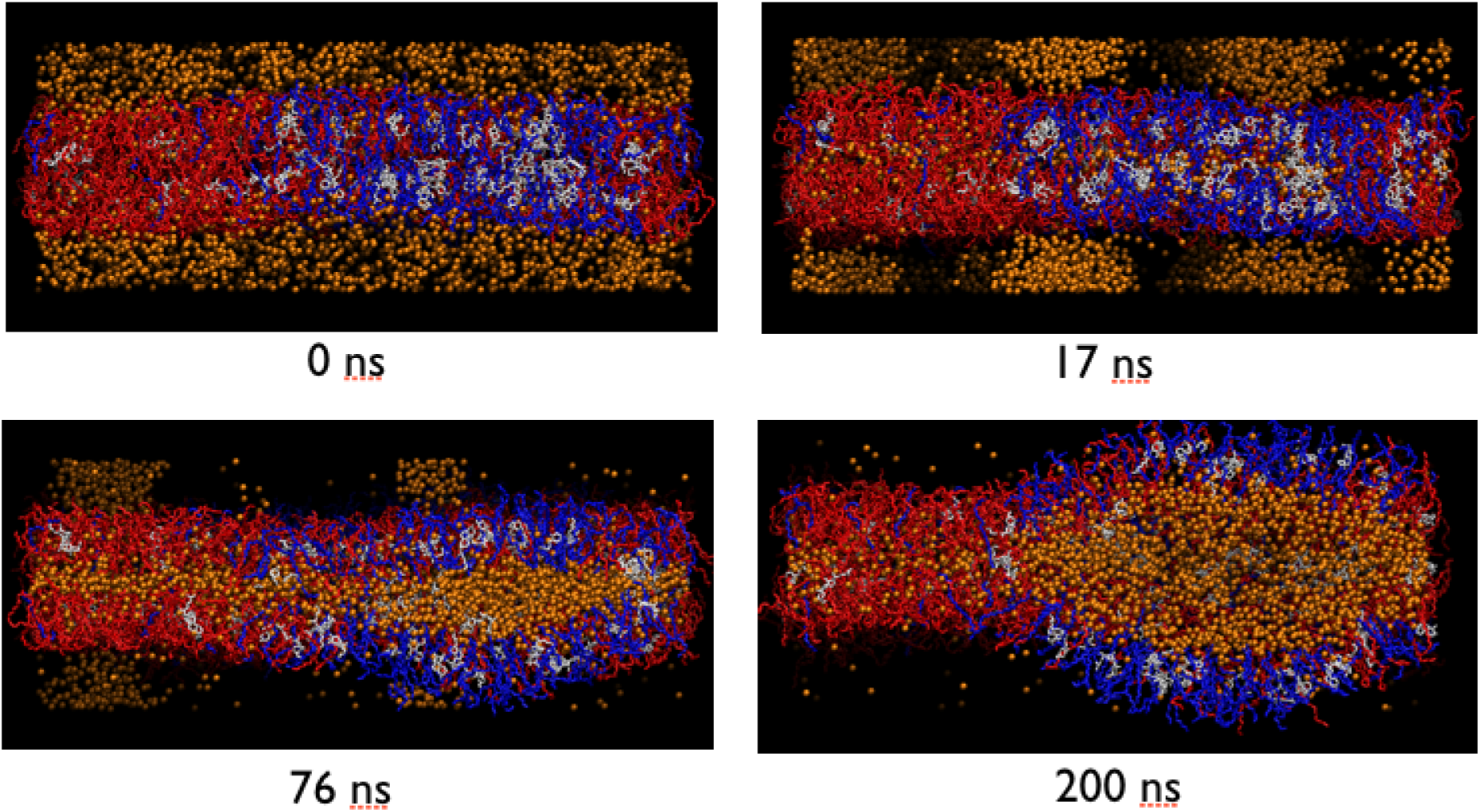
Nucleation and penetration of Xe nano-clusters in the membrane at a temperature of 323K.

